# CD9 marks myeloid/MegE-biased human hematopoiesis

**DOI:** 10.1101/2023.09.06.556527

**Authors:** Fatemeh Safi, Parashar Dhapola, Mikael N.E. Sommarin, Göran Karlsson

**Author notes:** Correspondence: Göran Karlsson, BMC B12, 22184 Lund, Sweden.

## Abstract

Rare hematopoietic stem cells make up an infrequent but critical population in the bone marrow (BM), maintaining and replenishing the entire hematopoietic system. Importantly, despite sharing the unique stem cell properties of multilineage differentiation and self-renewal, individual HSCs are functionally heterogeneous, and this heterogeneity increases during aging. While HSCs in young mice are qualitatively more similar, ageing is marked by an increased size of the HSC pool and substantial functional variation of individual HSCs. CD9 is a cell surface marker that is highly expressed in HSCs in mice, while CD9 expression within the human HSC population has been reported to be low during neonatal hematopoiesis. Here, we have investigated CD9 expression levels in the human HSPC population over time and identified that early in life; CD9 is infrequent in HSCs, but marks progenitor populations with low engraftment potential and high proliferation capacity. However, during situations of myeloid/Megakaryocyte-erythoid (MegE) biased hematopoiesis, such as during ageing or in leukemia, there is a substantial increase of CD9 expression in HSPCs. Thus, CD9 represents an HSC marker for myeloid/MegE-biased hematopoiesis.

## Introduction

While self-renewal capacity and differentiation to all blood lineages are the hallmarks of all hematopoietic stem cells (HSCs) ^1^, the stem cell pool is a heterogeneous population in multiple aspects, including the degree of self-renewal capacity ^2, 3^ as well as differentiation manner ^4-6^ and lifespan ^6-9^. Single-cell transplantation and in vivo fate mapping have shown that HSCs can contribute selectively to hematopoietic lineages with stable myeloid- or lymphoid-biased HSC subsets existing already in young mice ^10-12^.

Aging is linked to a number of changes in the human hematopoietic system leading to numerous adverse effects on hematopoiesis and immunity, as well as increased risk of certain hematopoietic diseases, including anemia ^13^, myelodysplastic syndrome (MDS), myelopoliferative disorders (MPDs) and myeloid leukemia ^14^, as well as reduced adaptive immune response ^15, 16^. These age-associated changes of hematopoiesis additionally correlate with an increase in immunophenotypic HSCs with decreased regenerative potential and engraftment capacity ^17-20^, as well as myeloid/platelet biased lineage differentiation ^17, 18, 21^, and up-regulation of MegE-biased gene expression programs ^22^. A recent comprehensive single cell transcriptomics analysis confirms more heterogeneity in the murine aged HSCs pool compared to the young and reveals the robust HSC transcriptomic aging signature ^23^.

Multiple sorting strategies have been developed to purify functionally different murine primitive HSCs based on cell surface markers heterogeneously expressed within the HSC population. For example, lymphoid-biased HSCs with high self-renew capacity have been shown to be marked by CD86 ^24^, and this marker is down regulated with aging ^25^. In contrast, myeloid-biased HSCs (CD150^+^)^26^ and platelet-biased HSCs (vWF) ^27^ with high self-renewal capacity are up regulated with aging ^21, 25^. It has also been reported that myeloid-biased HSCs with low self-renewal capacity (CD41^+^) are increased in frequency during aging of mice ^28^. Alteration in the expression of these markers correlate with functional activity and changes in heterogeneity of aged HSCs, for example the upregulation of CD150 and downregulation of CD86 is consistent with the myeloid–biased phenotype of aged HSCs ^29^.

In contrast to mouse, immunophenotypic changes associated with ageing effects are poorly explored in human hematopoiesis. Thus, further dissecting also the human HSC population has the potential to improve our understanding for the role of HSCs function and heterogeneity during aging and could possibly have an impact for treatment for leukemia or other age-related hematopoietic malignancies.

We have previously identified the tetraspanin CD9 as a marker that captures all murine HSCs ^30^, as well as multipotentiality during the earliest stages of hematopoietic lineage commitment ^31^. In human hematopoiesis, CD9 is expressed in stromal cells and influence physical interaction that has been suggested to regulate the differentiation of hematopoietic stem cells and progenitor in to megakaryocytes, B lymphoid and myeloid lineages (Brosseau et al. 2018). CD9 has also been described as a cancer stem cell (CSC) marker in diseases such as B acute lymphoblastic leukemia and acute myeloid leukemia (AML)^32-34^. Moreover, CD9 is expressed in 40% of human AML samples and associates with a favorable clinical outcome ^35^. In line with this, CD9 has been reported to be upregulated in AML leukemic stem cell (LSC) populations and is as such a potential AML stem cell marker ^34^. However very little is known regarding the role of CD9 as a marker for human HSCs.

Here, we thoroughly investigate the potential for CD9 as a marker for human HSCs. By combining flow cytometry analysis and CITE-seq as well as single-cell qPCR analysis and transplantation assays we identify a shift in CD9 expression during aging. In neonatal life CD9^+^ HSPCs display a myeloid and megakaryocytic lineage program with low engraftment potential and high proliferation capacity, while CD9^-^ cells are lymphoid primed and readily reconstitutes human hematopoiesis in xenograft models. CD9 expression is also altered during life from being primarily observed in MPPs and LMPPs, to be present in HSCs at high proportion in aged BM. This change of CD9 expression correlates with the age-dependent differentiation bias towards myeloid and MegE lineages and is especially apparent after leukemic transformation. Taken together, our data suggest that in human, CD9 expression in HSCs marks myeloid/MegE-biased hematopoiesis. ^36^

## Results

### The proportion of CD9+ HSPCs increased significantly with age

Even though it has been suggested that CD9 marks cancer stem cells in B acute lymphoblastic leukemia (B-ALL) as well as in acute myeloid leukemia (AML) ^32-34^, very little is known regarding the role of CD9 in healthy human HSCs. Using flow cytometry analysis we investigated CD9 expression in the LIN^-^CD34^+^CD38^-^ HSPCs population from neonatal umbilical cord blood (CB) as well as in young Bone Marrow (yBM, 20-30 years old) and aged Bone Marrow (aBM, 50-63 years old). In accordance with previous studies only a small fraction of CB and yBM HSPCs expressed cell-surface CD9 ^34^ while the proportion of CD9^+^HSPCs substantially expanded with age (Fig-1A, B). To investigate if this increase is due to a general upregulation of CD9 across different HSPC specific sub-populations, the HSPCs were divided using established cell surface markers for lympho-myleoid progenitors (CD45RA) or for HSCs (CD90) (FIG. 1C, D). We observed that CD9^+^ cells are primarily detected in multipotent progenitors (MPPs), and multi-lymphoid progenitors (MLPs/ LMPPs) with low expression in the HSCs population in CB and yBM (FIG. 1E). With age there was a shift in this HSPC distribution where LMPPs were lost in combination with a relative increase of the MPP population, importantly containing a larger fraction of CD9^+^ cells (FIG. 1C-F).

**Figure 1.**
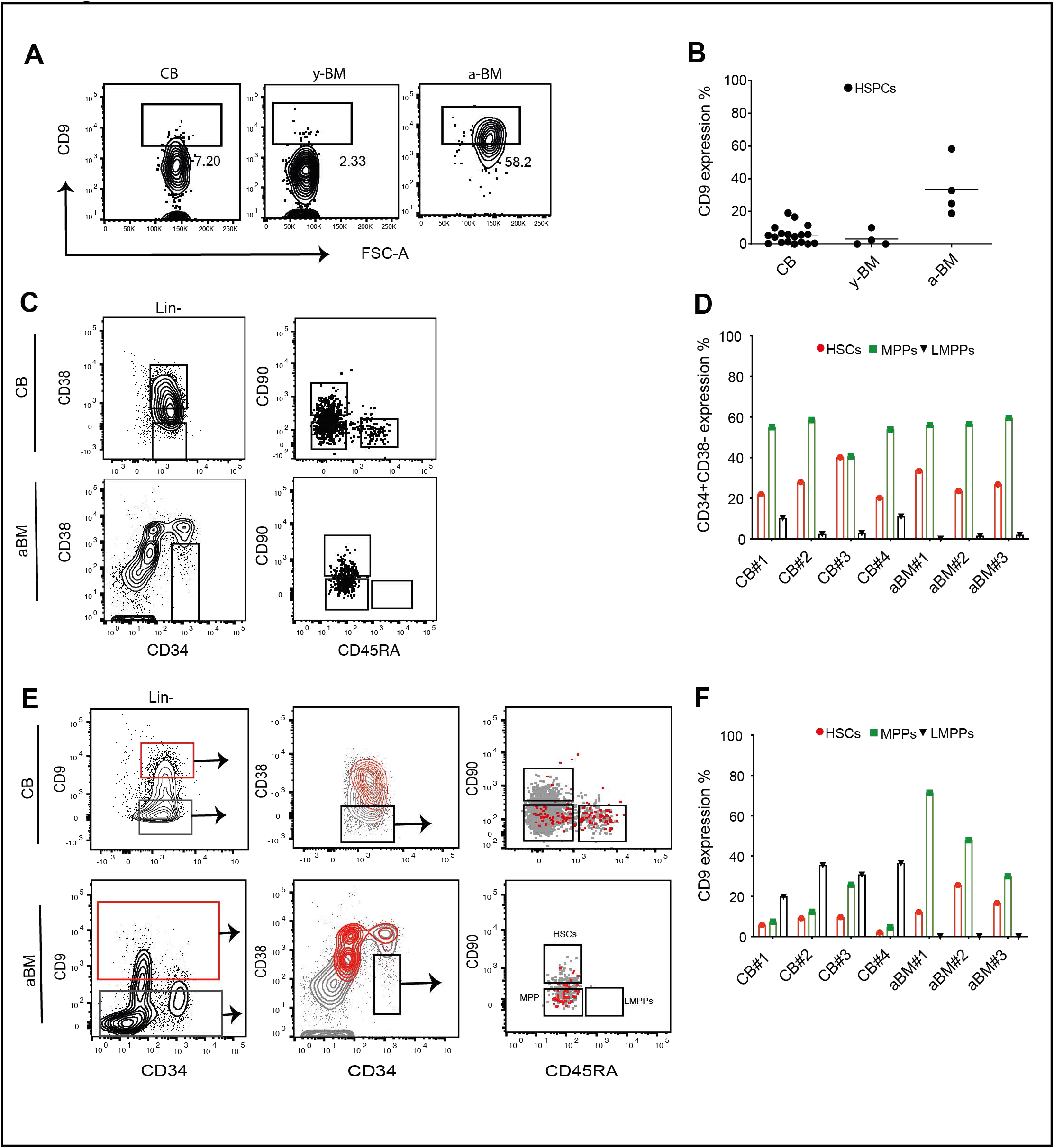
The CD9^+^ HSCs population and CD9^+^ LMPP are expanded and decrease respectively in old people. **A)** Representative FACS plots for CD9 expression in the HSPCs compartment (LIN-CD34^+^CD38^-^) in CB, yBM and aBM samples. **B)** Pooled data for CD9 expression in the LIN^-^CD34^+^CD38^-^ population in (CB n=18, young BM n=4, aged BM n=4). **C)** Representing FACS plots for analysis of HSPCs progenitor populations and **(D)** compiled data for CB n=4 and aged BM n=3. **E)** Representative FACS plots for CD9 expression in CD34^+^CD38^-^ HSCs (CD90^+^CD45RA^-^), MPP (CD90^-^CD45RA^-^) and MLPs (CD90^-^CD45RA^+^) and compiled data for CB n=4 and aged BM n=3 **(F)**

### CD9 expression increases as HSPCs heterogeneity changes towards HSCs and MegE differentiation during ageing

To directly compare the CD9 expression in HSPCs across the different age groups and further correlate CD9 expression with changes in heterogeneity, we used our in house generated single-cell CITE-seq data on from CD34^+^CD38^-^ cells of CB, yBM, and aBM ^37^. Data from all age groups were combined and visualized using a two dimensional UMAP (FIG. 2A), followed by clustering identifying seven subpopulations based on distinct molecular signatures (FIG. 2B). To track the differentiation axis, pseudotime values were calculated for the cells and visualized on the UMAP (FIG. 2C). Here, cluster 1 contains the most primitive cells and is followed by two trajectories of stepwise differentiation into cluster 5 and 6. Each cluster was annotated based on expression of cell type-specific marker genes as defined by available gene expression databases for various hematopoietic populations^38^.The cell type specification for each cluster was visualized by radar plots (FIG. 2D). As expected, Cluster 1-exhibited enrichment in cells with an HSC molecular signature, while cluster 2- and 6-contained myeloid/MegE, and MegE-committed signatures, respectively. In contrast cluster 3 displayed an LMPP/lympho-myeloid gene expression program with increasingly lymphoid-committed signatures in cluster 4 and 5. Interestingly, the proportion of stem cell and myeloid/MegE-biased clusters within the HSPCs dramatically increased with age, while the lympho-myeloid clusters decreased (FIG. 2E). These observations within the most primitive HSPCs are inline with known ageing effects on hematopoiesis, from lympho-biased at birth to myeloid and MegE-biased in the elderly ^22, 39, 40^. CITEseq allows for directly combining single-cell gene expression analysis with high throughput immunophenotypic profiling. Visualizing CD9 cell surface expression (CD9-ADT) in the UMAP revealed that CD9 is mainly expressed within the HSC and myeloid/MegE-biased clusters (FIG. 2F, G). Thus, CD9 expression divides even the earliest hematopoietic cells and correlates with populations known to have myeloid/MegE potential. This is in line with studies showing that CD9 is upregulated during differentiation of MegE progenitors ^41^. Moreover, when comparing the number of CD9^+^ and CD9^-^ cells in each cluster during aging, CD9 was found to be highly expressed in the LMPP/lympho-myeloid cluster 3 in CB, while the lymphoid-biased cluster 4 was marked by a lack of CD9 expression. Interestingly, these two clusters were both substantially proportionally decreased during ageing. In contrast the CD9 rich HSC cluster 1 and myeloid/MegE-biased clusters 2 and 6 in CB were proportionally increased during ageing (FIG. 2H, I).

**Figure 2.**
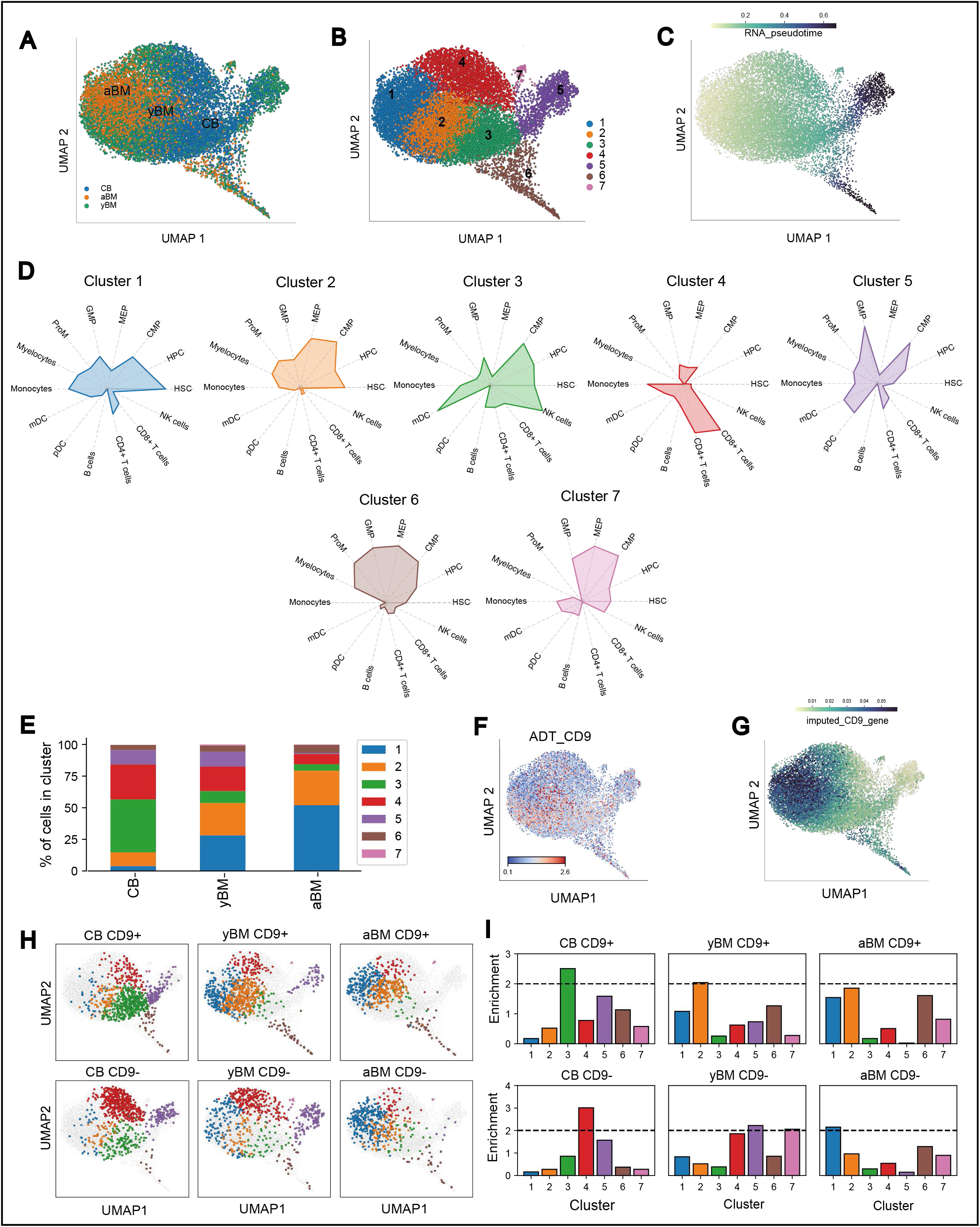
During aging HSPCs heterogeneity changed toward HSCs and Meg differentiation with an increase of CD9 expression. **A)** UMAP visualization of merged yBM, aBM, and CB derived 14,010 CD34^+^CD38^-^ cells. **B)** UMAP showing the five clusters identified from the merged data. **C)** UMAP showing pseudotime scores derived using HSCs cluster as source cluster. **D)** Radar plots showing cell type signature of maker genes from each of the clusters. **E)** Percentage of total cells in each cluster. **F)** Normalized values of CD9 ADT, red marks high expression though blue marks low expression **G)** UMAP showing imputed expression values of CD9 gene. **H)** UMAP showing the cells identified as either CD9^+^ or CD9^-^ in each of the sample source. **I)** Bar graphs showing the enrichment of CD9^+^ or CD9^-^ cells in each of the clusters across sample sources.

To further define the CD9^+^ cells within the most primitive LIN^-^CD34^+^CD38^-^ HSC population we performed single-cell multiplexed quantitative reverse transcription (qRT)-PCR analysis on a microfluidics–based platform. 160 CD9^+^ and 125 CD9^-^ cells were run against a pre-selected set of 96 gene-primers including genes of functional HSC regulators, as well as genes involved in lineage commitment, and cell cycle (FIG. 3.A). Following principal component analysis (PCA), we observed that CD9^+^ and CD9^-^ cells express distinct molecular signatures in CB with little overlap between them (FIG. 3.B). CD9^-^ cells contained a molecular signature associated with lymphoid commitment such as *CD10, CD11a, Sterile-Igh, IRF8* and *RAG*, which was only found in neonatal cells. In contrast CD9^+^ cells of any age expressed genes associated with HSC function like *MECOM* and ETS2, as well as with myeloid/MegE-related genes like *GATA2, KIT, FOXO1* and *TAL1* (FIG. 3.C). Interestingly aged CD9^-^ HSCs express a similar gene signature as compared to CD9^+^ cells including megakaryocyte genes like GFI1B and CD41 (FIG. 3.D). This data together with the CiteSeq analysis suggest that CD9 marks myeloid/MegE biased hematopoiesis. This is in agreement with the observed reduction of CD9^-^ HSCs, and expansion of CD9^+^ HSCs during myeloid-biased hematopoiesis associated with age and myeloid leukemia.

**Figure 3.**
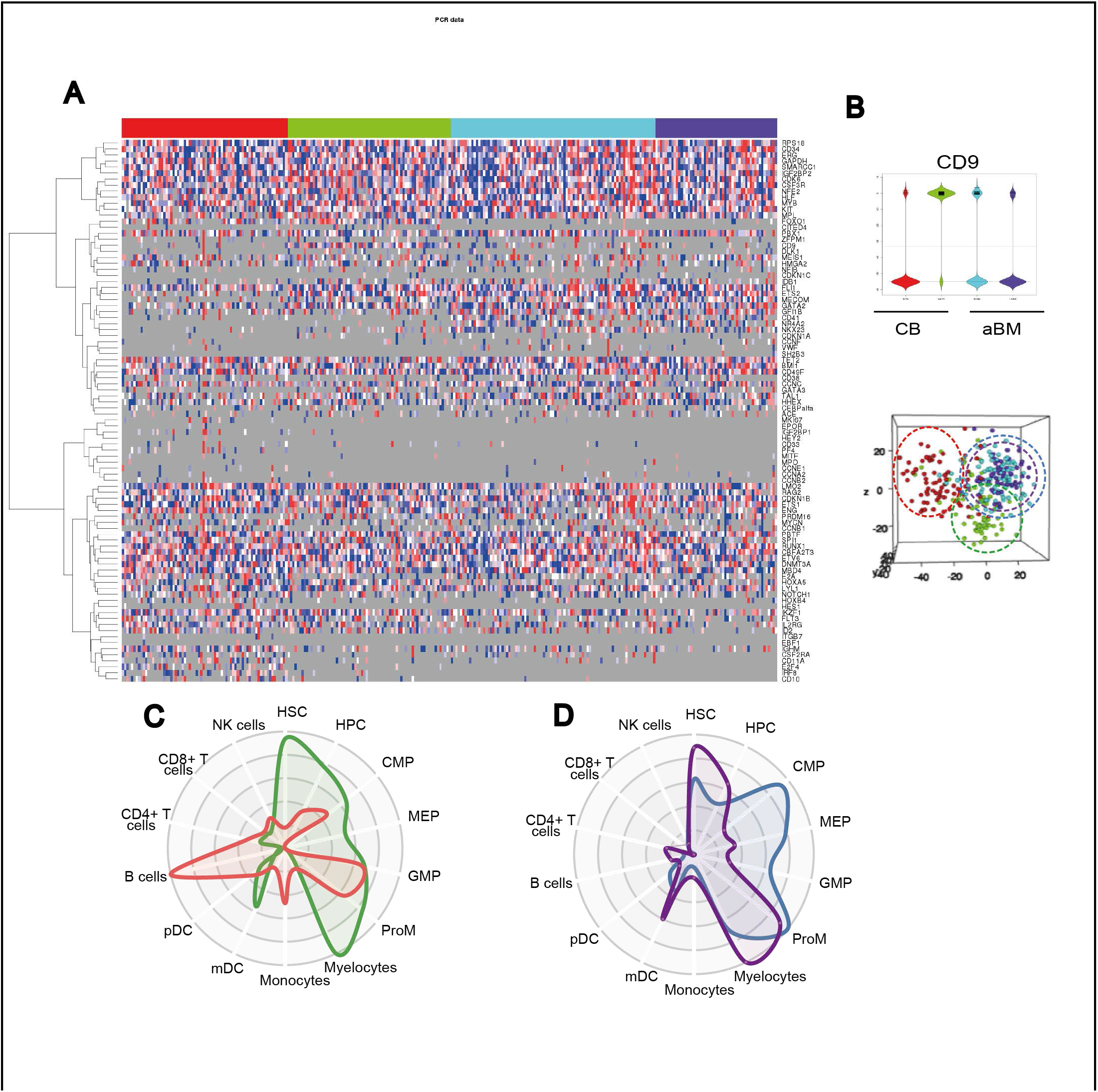
CD9 discriminates between HSC/My/Ery and lymphoid molecular programs within the human HSPC population. **A)** Fluidigm single-cell gene comparing expression of 96 genes across 160 CD9^+^ and 125 CD9^-^ LIN^-^ CD34^+^CD38^-^ single cells in CB or old BM. **B)** Unsupervised principle component analysis (PCA) reveal that CD9^+^ and CD9^-^ CB HSPCs express distinct molecular programs, **C)** Radar plot of the gene with the most variable expression demonstrating that CD9^-^ CB HSPCs (in red) represent a subpopulation expressing a lymphoid molecular program that is lost with age (purple) and CD9 high in CB HSPCs (green) gain with age (blue) a subpopulation with expressing MEP and GMP signature.

In order to compare HSCs activity between CD9^+^ and CD9^-^ HSCs, 100 CD9^+^ or CD9^-^ LIN^-^ CD34^+^CD38^-^ cells were sorted from CB and kept in culture under minimum cytokine conditions for 14 days. Cell counting revealed that CD9^+^ cells were more proliferative compared to CD9^-^ cells (FIG. 4.A) This finding was supported by single-cell proliferation experiments where CD9^+^ HSCs entered cell cycle significantly faster as compared to CD9^-^ HSCs. Following 66 hours of culture 2.6 fold more CD9^+^ cells had divided compared to CD9^-^ (64% CD9^+^ vs. 24% CD9^-^) cells reached 76% division frequency 18 hours later than CD9^+^ cells (90h compare to 72 hours) (FIG. 4.B).

**Figure 4.**
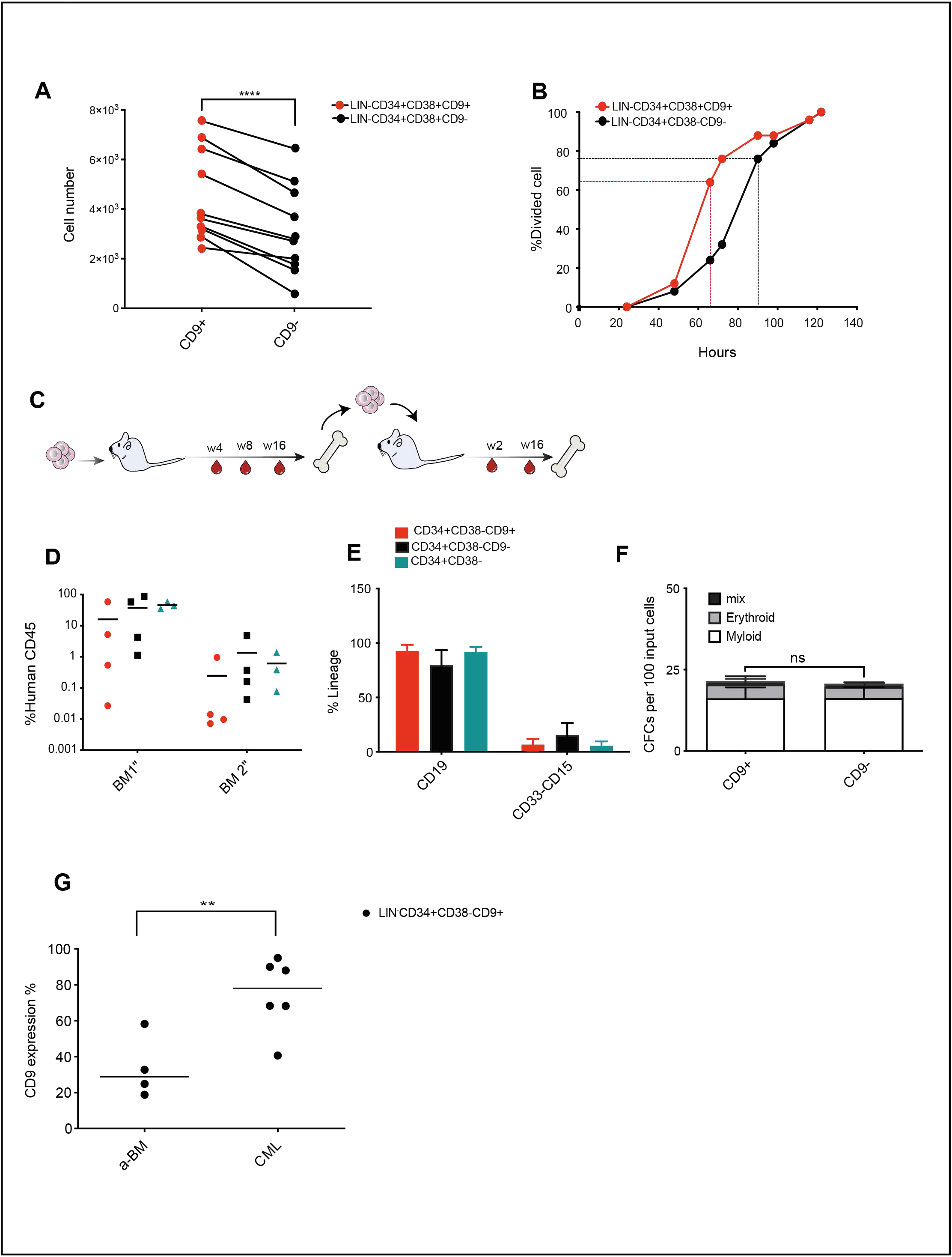
Both the CD9^+^ and CD9^-^ h-HSPC populations contain in vivo repopulation activity, however CD9^+^ HSPCs more rapidly enter cell cycle. **(A)** Growth ability of CD9^+^ cells and CD9^-^ cells between 10 individual across the 4 biological replicate of CB, ****p<0.0001. **(B)** Cumulative first division kinetics (excluding dead cells) of CD9^+^ (red) and CD9^-^ (black) in single-cell HSPCs (>80 single cells). (**C)** Experimental design for transplantation of 1000 CD9^+^ or CD9^-^ LIN^-^CD34^+^CD38^-^ cells into sub-lethally irradiated NSG mice. (**D)** Human reconstitution in the BM of recipient mice following 16 weeks of primary and secondary transplantations and (**E)** the mean lineage distribution in the BM following primary transplantation (n= 4 mice in 2 individual experiments). **(F)** Number of CFCs per 100 input cells (n= 3 individual experiments, not significant) **(G)** CD9 expression in the CD34^+^CD38^-^ of CML (Diagnosis n= 6) compared to old BM from figure 1, **p<0.01.

To further characterize the functionality of CD9^+^ and CD9^-^ LIN^-^CD34^+^CD38^-^ cells in CB, we performed xenotransplantation experiments into immunodeficient NSG mice (FIG. 4C). However, despite the differences observed in the molecular analysis, both CD9^+^ and CD9^-^ CB HSCs displayed similar long-term engraftment potential and lineage differentiation capacity (FIG. 4D, E).

Similarly, when performing CFC assays where 100 HSCs were cultured for 14 days in semisolid methylcellulose medium supplemented with cytokines supporting myeloid and erythroid differentiation, there was no significant difference in colony forming capacity between the CD9^+^ and CD9^-^ HSPC populations (FIG. 4F).

Thus, standard in vitro and in vivo methods revealed no significant differences between CD9^+^ and CD9^-^ cells when it comes to primitive HSPC function, suggesting that CD9 expression is rather an indicator of molecular priming towards certain lineages. To further confirm the potential for CD9 to mark lineage bias we performed FACS analysis on Chronic Myeloid Leukemia (CML) patient material. The BCR-ABL mutation associated with CML is known to bias early HSC differentiation towards myeloid and megakaryocytic lineages ^42^. Therefore, CML is a good model to measure CD9 expression in HSCs with known myeloid/MegE bias. Indeed, CD9 was highly expressed as measured by FACS analysis of HSCs from all CML patients analyzed (Figure.4H), and on average 2.2 fold more CML HSPCs expressed cell-surface CD9 compared to age matched healthy controls (FIG. 1) (p<0.01).

## Discussion

Here, we explored the capacity for CD9 as a marker for distinct cell types in human HSPCs. Our data indicate that while CD9 may not differentiate functionally distinct cells, its expression, even in the most primitive HSPCs, correlates with downstream myeloid/MegE-biased hematopoiesis, both in health and disease. This suggests that CD9 expression is an indicator of molecular priming towards myeloid/MegE commitment. To investigate whether CD9 expression signifies priming of HSPCs towards myeloid or MegE cells, particularly during aging, applying ATAC-seq analysis and motif identification in enhancer regions ^43, 44^ would be intriguing. As the enhancer landscape provides a more accurate reflection of cell identity compared to mRNA levels ^45^. An alternative approach for directly measuring clonal lineage contribution involves using cell barcoding to trace cell fate and assess lineage commitment. The scATAC-seq technique offers an elegant solution by utilizing reads mapped to the mitochondrial genome for clonal lineage tracking while simultaneously capturing chromatin features within the same cell ^46^. Novel technology developing continues which led to new understanding from hematopoiesis system.

## Methods

### Sample preparation

Human cord blood (CB) was collected from umbilical cords at the end of full deliveries obtained from the Obstetrics departments in Lund or Malmo, Sweden, with the mother’s informed consent. Human bone marrow (BM) aspirates were obtained from informed donors from the department of Haematology in Lund, Sweden BM from CML patients were aspirated from the posterior iliac crest after informed consent according to protocols approved by the regional research ethics committees of sites in Lund, Helsinki, Uppsala, Aarhus, and Stockholm. All samples were enriched for mononuclear or CD34^+^ cells and cryopreserved prior to analysis.

Mononuclear cells were isolated from CB and BM on Ficoll gradient according to guidelines approved by Lund University. CD34 enrichment was preformed using CD34 enrichment kit (Miltenyi Biotec, Bergisch Gladbach, Germany) as previously describe ^47^ in brief, cells were stained with FC block and CD34 enrichment beads for 20 min, then positive magnetic enrichment was performed on the column. After either MNC or CD34 enrichment all cells were frozen in FBS with 10% DMSO and kept at -150°C until use.

### Flow cytometry analysis and cell sorting

CD34^+^ CB or BM was stained with fluorophore-conjugated human antibodies in PBS with 2% Fetal Bovine serum (FBS) (Hyclone), lineage cocktail (CD14 (eBiocience), CD16 (Biolegend), CD19 (Biolegend), CD56 (Biolegend), CD2 (Biolegend), CD3 (Biolegend), CD123 (Biolegend), and CD235a (BD Biocience), CD34-FITC (Biolegend), CD38-APC (Biolegend), CD45RA-PB (BD Biocience), CD90-BV605 (BD Biocience) and CD49F-PECY7 (eBiocience), CD9-PE (eBiocience) for 30 min in 4°C. Dead cells were excluded with 7-aminoactinomycin D (7-AAD, Sigma). Cell were either sorted on AriaII or AriaIII cell sorter (BD Biosciences, San Jose, CA) with a purity >95% into sterile filtered PBS with 2% FBS or acquired on a FACS Canto II or LSR fortes Analyzer (BD Biosciences). Single cells were index sorted using single cell depositor. Flow cytometer data was analyzed with FlowJo software (v10, Tree Star Inc)

### CITE-seq data analysis

#### UMAP and clustering of the data

The CD34^+^CD38^-^ populations from yBM (4,882), aBM, (3,967) and CB (5,161) samples (total 14,010 cells) from ^37^ were merged using the partial learning approach of Scarf ^48^. The neighborhood graph of cells was constructed using top 21 principal components and 15 neighbours. UMAP embedding for this graph was calculated as implemented in Scarf (with 2000 training epochs). Clustering was performed using Leiden algorithm with resolution parameter set at 0.6. The pseudotime values were calculated using Scarf’s ‘run_pseudotime_scoring’ function (which internally uses PBA algorithm, ^36^) setting cluster 1 as source and cluster 5, 6 and 7 as sinks.

#### Marker identification

Scarf’s ‘run_maker_search’ function was used to identify genes that were associated with each of the clusters. The cells from sample yBM samples were used to identify the genes that are anti/correlated with imputed CD9 expression (‘get_imputed’ function of Scarf that uses MAGIC algorithm ^49^. For CellRadar visualization, the data from BloodSpot ‘normal human hematopoiesis’ was used ^38^ to annotate cell type identity of each cluster. The values for the radar plot were generated using min-max scaled median value of input genes.

#### Identification of CD9^+^ and CD9^-^ cells

To identify the CD9^+^ and CD9^-^ genes in each sample (CB, yBM, and aBM) z-scores of CLR normalized CD ADT values were used. The cells with zscore > 1 were considered CD9^+^ while cells with zscore < -1 were considered CD9^-^. Enrichment of CD9^+^ cells in each cluster within each sample was calculated as:

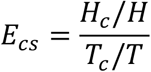

Where *E*_*CS*_ is the enrichment of CD9^+^ cells in a cluster *c* in sample *s. H*_*C*_ is the number of CD9^+^ cells in a given cluster, *H* is the total number of CD9^+^ cells in a given sample, *T*_*C*_ is the total number of cells in a the same cluster, *T* is the total number of cells in the sample.

Number of cells in each group

CB: CD9^+^: 737 CD9^-^: 766

yBM: CD9^+^: 773 CD9^-^: 619

aBM : CD9^+^: 519 CD9^-^: 474

### Sc-qPCR and data analysis

Multiplexed quantitative real-time PCR analyses (BioMark) 96.96 Dynamic Array platform (Fluidigm) with Taqman Gene Expression Assays listed in the supplemental table 1 (Applied Biosystems, CA, USA) were performed on index-sorted cells from LIN^-^D34^+^CD38^-^CD9^+^/CD9^-^ population as described previously ^50^. In Brief, single cells were index-sorted on 96 well plates containing 4ul lyses buffer with 0.4%NP40, deoxynucleoside triphosshates (dNTPs), dithiothreitol (DTT) and RNasOUT (Invitrogen), and snap frozen. Pre-amplification was performed using Taq-Man primers and Taq/SSIII reaction mix (Invitrogen), according to prep program; for 25 cycle and diluted 5 times prior to real-time q-PCR. cDNA was added to 96.96 dynamic Array Chips (Fluidigm) according to manufacturer’s instructions with the individual Taq-Man assays. The data analyzed using the SCExV tool ^51^.

### Xenotransplantation and in-vivo

All mice studies were performed at Lund University and were approved by the local ethical committee. 1000 CB cells (LIN^-^ CD34^+^ CD38^-^ CD9^+^ or CD9^-^ and control positive LIN^-^ CD34^+^ CD38^-^) were transplanted intravenously in 8 to 12 week old sub-lethally irradiated (250 cGY) NOD.Cg-Prkdc^scid^ Il2rg^tm1Wjl^/SzJ (NSG;Jackson laboratories) mice. Contribution of human cells in Peripheral blood (PB) was measured every 4-week, using mice antibody for CD45-FITC and human antibodies for CD45-APC, CD15-PE, CD33-PE and CD19-BV605. 16 week following transplantation Tibia and femur were collected from both legs and the BM was stained with mouse CD45-AF700, human antibodies CD45-APC, CD15-PE, CD33-PE, CD34-FITC and CD19-BV605, all PB and BM was analysed for human engraftment on an LSR Fortessa (BD). For secondary transplantation, we injected a half-tibia/femur equivalent of BM cells from primary transplanted mice to the new irradiated recipients mice.

### In vitro culture and progenitor cell assays

For cell growth assay 100 LIN^-^CD34^+^CD38^-^ CD9^+^ or CD9^-^ cells were sorted and seeded in 96 well plate and culture in StemSpan Serum –Free Expansion Medium (SFEM, StemCell Technologies) supplemented with 100u/ml penicillin (Hyclone) and 0.1 mg/ml streptomycin (hyclon) (PEN/STREP (1%)) and SCF, TPO, FLT3L from Peprotech that used at (100ng/ml), after 2 week incubation in 37°C, cell were counted with Neubauer chamber slide. For time first division Single cell LIN^-^CD34^+^ CD38^-^ CD9^+^ or CD9^-^ cells sorted in to the round bottomed 96 well plates and culture in SFEM (StemCell Technologies), supplemented with SCF (10ng/ml), TPO (10ng/ml), Flt3L (2ng/ml), g-CSF (10ng/ml), IL6 (2ng/ml), IL3 (2ng/ml) from Peprotech and EPO (2U/ml). Colony forming cell (CFC) assay done by culturing 100-sorted cells LIN^-^CD34^+^ CD38^-^ CD9^+^ or CD9^-^ in to MethoCult™ H4434 with human cytokine (Stem Cell Technologies) and colony formation score at day 12-14 of incubation in 37°C.

### Statistical analysis

Statistical significance performed in GraphPad Prism (v.8.2) using paired student test, error bar represent standard deviation (SD).

## Acknowledgements

We express our gratitude to Johan Richter, MD, PhD, for providing the bone marrow (BM) samples, as well as to all the donors who contributed cord blood and BM samples. Additionally, we thank Anna Fossum and Zhi Ma at the FACS Core facility at Lund Stem Cell Center for their expert support in flow cytometry. We would like to acknowledge Clinical Genomics Lund, SciLifeLab and Center for Translational Genomics (CTG), Lund University, for providing expertise and service with sequencing and analysis. This work was supported by grants from the Swedish Cancer Society, the Ragnar Söderberg Foundation, the Knut and Alice Wallenberg Foundation, the Swedish Research Council, the Swedish Society for Medical Research, and the Swedish Childhood Cancer Foundation.

## Author contributions

G.K. F.S. Conceived and designed the study; F.S. designed and performed the experiments; P.D. designed and performed the bioinformatics analyses. M.S. provided the single-cell CITE-seq data; F.S. and G.K. analyzed and interpreted data; F.S. wrote the manuscript and prepared the figures with contribution from GK; GK supervised the study; all authors reviewed, edit and approved the manuscript.

## Data availability

The datasets generated and analyzed during the current study are available from the corresponding author on reasonable request. Additionally, the single-cell CITE-seq data used in this study were obtained from a previously published dataset ^37^.

## Supplementary information

Supplementary Table 1 contains the list of Taqman Gene Expression Assays that used in Fluidigm experiment.

